# Assessing circular RNAs in Alzheimer disease

**DOI:** 10.1101/800342

**Authors:** Laura Cervera-Carles, Oriol Dols-Icardo, Alba Cervantes-Gonzalez, Laura Molina-Porcel, Laia Muñoz-Llahuna, Jordi Clarimon

## Abstract

Circular RNAs (circRNA) are evolutionary conserved non-coding RNAs resulting from the backsplicing of precursor messengers. Recently, a circular-transcriptome-wide study of circRNA in brain tissue from patients with Alzheimer’s disease (AD) has revealed a striking association between the expression of circRNA and AD pathological diagnosis. In the present study, we aimed at replicating the major findings in an independent case series comprising definitive sporadic and familial AD. In order to assess the specificity of circRNA changes, we also included cases with frontotemporal lobar degeneration (FTLD), comprising brain specimens with TDP-43 aggregates (FTLD-TDP43) and samples that presented Tau accumulation (FTLD-Tau). Through a quantitative PCR approach, we evaluated a total of eight circRNAs that surpassed the significant threshold in the former meta-analysis (*circHOMER1*, *circDOCK1, circKCNN2*, *circMAN2A1, circFMN1, circRTN4, circMAP7*, and *circPICALM).* Average expression changes between AD patients and controls followed the same directions as previously reported, suggesting an overall upregulation of *circDOCK1*, *circMAP7*, *circMAN2A1*, *circRTN4* and *circPICALM*, and a downregulation of the remainder (*circHOMER1, circFMN1* and *circKCNN2*) in AD brain tissue. We also confirmed an exacerbated alteration in circRNA expression in the Mendelian AD group compared to the sporadic forms. Two circRNAs, *circHOMER1* and *circKCNN2*, also showed significant expression alterations in the group of FTLD-Tau and FTLD-TDP43, respectively. Overall, these results reinforce the conception that expression of circRNAs is altered in Alzheimer’s disease, and also suggest a wider involvement of this particular class of RNA in other neurodegenerative dementias.

## Introduction

Circular RNAs (circRNAs) are a class of low-abundance biochemically stable single stranded RNAs, produced by the back-splicing of exons in precursor mRNAs. Although their role in the central nervous system is poorly understood, the enriched expression in brain tissue and their overall upregulation during neurogenesis and synaptic plasticity has fueled the interest of these noncoding family of transcripts to underscore the biological bases of neurodegenerative disorders^1–3^.

Recently, using deep RNA profiling in brain tissue from patients with Alzheimer disease (AD) and healthy controls, Dube *et al*. have shown evidence for changes in circRNA expression occurring early, in pre-symptomatic AD, as well as in Mendelian forms of the disease^4^. Through a circular transcriptome-wide analysis in two independent RNA-seq datasets, the authors disclosed more than 3.500 different circRNA species. Among them, 28 were significantly associated with the three traits that were investigated: neuropathological staging, dementia severity and differential expression between phenotypes. Correction for multiple comparisons disclosed a group of circRNAs that surpassed the p-value threshold for all three traits in their meta-analysis. In the present study we aimed at replicating some of the observed changes in an independent cohort of patients with a neuropathological diagnosis and assess whether these alterations are specific to AD.

## Methods

Total RNA was isolated from 60mg of frozen frontal cortex samples obtained from the Neurological Tissue Bank (Barcelona). Brain tissue was first homogenized with a mortar and pestle under liquid nitrogen and transferred into Trizol reagent (Life Technologies). The conventional phenol-chloroform protocol was followed by a column-based RNA purification method using the RNeasy Mini kit (Qiagen, Hilden, Germany). RNA integrity (RIN) was assessed using the 2100 Bioanalyzer instrument (Agilent). All RNA samples presented a RIN>6.

RNA was reverse transcribed with the RevertAid First Strand cDNA synthesis kit (Thermo Scientific, Rockford, IL). For quantitative polymerase chain reaction (qPCR), the obtained cDNAs were analyzed in duplicate in a 7900HT Fast Real-Time PCR system (Applied Biosystems, Foster City, CA), using the Fast SYBR Green Master mix (Thermo Scientific), 200 nM of each primer and 30 ng of cDNA. circRNAs were assayed using the previously reported primers^4^, and GAPDH was used as normalizer. Prior to the analysis a detection threshold for qPCR was established at 38 cycles. Expression levels were determined by the 2^−ΔΔCt^ method. Kruskal-Wallis non-parametric test with Dunn’s post-hoc analysis was applied for comparisons between diagnostic groups. Adjusted statistical significance was set at α=0.05.

Neuropathological examination was performed according to standardized protocols at the Neurological Tissue Bank of the Biobanc-Hospital Clinic-IDIBAPS^5^, and disease evaluation was performed according to international consensus criteria^6,7^.

## Results

In the present study we assessed the expression of 8 circRNA species that exceeded the stringent gene-based, Bonferroni multiple test correction in the final meta-analysis by Dube et al.: *circHOMER1*, *circDOCK1, circKCNN2*, *circMAN2A1, circFMN1, circRTN4, circMAP7*, and *circPICALM.* The frontal cortex of 69 brain samples was included in the analysis (Table 1). These included samples from patients with a definitive diagnosis of AD (n = 19), patients with frontotemporal lobar degeneration (FTLD) presenting aggregates of the transactive response DNA-binding protein 43 kDa (FTLD-TDP43, n=16), and patients with Tau pathology (FTLD-Tau, 6 with corticobasal degeneration and 4 with Pick’s disease, Table 1). Since individuals with autosomal dominant AD showed more dramatic changes in circRNA expression compared to sporadic AD in the Dube *et al*. study, we also added 9 brain specimens from familial AD patients harboring Mendelian mutations (fAD) (see Table 1 for details). Similarly, among the FTLD series 9 individuals harbored the hexanucleotide repeat expansion in the *C9ORF72* gene (a known genetic cause of FTLD-TDP) and 5 were carriers of the p.P301L missense mutation in the *MAPT* gene (leading to FTLD-Tau). Fifteen postmortem brain samples from neurologically healthy controls were also included. All samples were provided by the Neurological Tissue Bank at Hospital Clínic (IDIBAPS, Barcelona), and met the neuropathological diagnostic criteria for their respective diagnoses^6,7^. The study (IIBSP-DFT-2018–97) was approved by the local Ethics Committee of both the Tissue Bank and Sant Pau Research Institute.

**Table 1.** Demographic, clinical and genetic data of all brain specimens.

| Case | Diagnosis | Gender | Age at death (y) | PMI, (h:m) | Mutations |
| --- | --- | --- | --- | --- | --- |
| 1 | Control | Female | 68 | 13:00 |  |
| 2 | Control | Female | 81 | 23:30 |  |
| 3 | Control | Female | 83 | 07:30 |  |
| 4 | Control | Female | 83 | 07:30 |  |
| 5 | Control | Male | 64 | 10:00 |  |
| 6 | Control | Male | 78 | 06:00 |  |
| 7 | Control | Male | 31 | 17:30 |  |
| 8 | Control | Female | 50 | 12:00 |  |
| 9 | Control | Female | 37 | 13:30 |  |
| 10 | Control | Female | 74 | 03:40 |  |
| 11 | Control | Female | 56 | 14:00 |  |
| 12 | Control | Male | 71 | <24:00 |  |
| 13 | Control | Female | 75 | 20:00 |  |
| 14 | Control | Female | 78 | <24:00 |  |
| 15 | Control | Female | 77 | 20:00 |  |
| 16 | sAD | Male | 87 | 08:00 |  |
| 17 | sAD | Female | 83 | 20:00 |  |
| 18 | sAD | Male | 74 | 10:00 |  |
| 19 | sAD | Female | 74 | 14:33 |  |
| 20 | sAD | Female | 87 | 05:45 |  |
| 21 | sAD | Female | 75 | 04:00 |  |
| 22 | sAD | Female | 77 | 05:30 |  |
| 23 | sAD | Female | 83 | 10:00 |  |
| 24 | sAD | Female | 77 | 04:30 |  |
| 25 | sAD | Female | 86 | 09:00 |  |
| 26 | sAD | Male | 83 | 05:00 |  |
| 27 | sAD | Male | 75 | 04:15 |  |
| 28 | sAD | Female | 80 | 05:30 |  |
| 29 | sAD | Female | 73 | 14:00 |  |
| 30 | sAD | Female | 83 | 04:30 |  |
| 31 | sAD | Female | 61 | 04:30 |  |
| 32 | sAD | Female | 66 | 07:00 |  |
| 33 | sAD | Male | 56 | 16:45 |  |
| 34 | sAD | Male | 68 | 09:00 |  |
| 35 | fAD | Male | 57 | 15:15 | *PSEN1* (p.M139T) |
| 36 | fAD | Male | 44 | 05:30 | *PSEN1* (p.E120G) |
| 37 | fAD | Male | 57 | 09:30 | *PSEN1* (p.V89L) |
| 38 | fAD | Female | 56 | 05:00 | *PSEN1* (p.L286P) |
| 39 | fAD | Female | 56 | 06:00 | *PSEN1* (p.P264L) |
| 40 | fAD | Male | 64 | 14.45 | *PSEN1* (p.M139T) |
| 41 | fAD | Male | 36 | 15:00 | *APP* (p.I716F) |
| 42 | fAD | Male | 53 | 05:15 | *PSEN1* (p.M139T) |
| 43 | fAD | Male | 68 | 06:10 | *APP* (duplication) |
| 44 | FTLD-TDP43 | Female | 70 | 11:00 |  |
| 45 | FTLD-TDP43 | Male | 72 | 17:30 |  |
| 46 | FTLD-TDP43 | Female | 69 | 11:45 |  |
| 47 | FTLD-TDP43 | Male | 65 | 13:00 |  |
| 48 | FTLD-TDP43 | Male | 77 | 06:20 |  |
| 49 | FTLD-TDP43 | Male | 73 | 13:00 |  |
| 50 | FTLD-TDP43 | Male | 71 | 07:00 |  |
| 51 | FTLD-TDP43 | Male | 61 | 07:45 | *C9orf72* |
| 52 | FTLD-TDP43 | Male | 66 | 15:15 | *C9orf72* |
| 53 | FTLD-TDP43 | Male | 69 | 05:00 | *C9orf72* |
| 54 | FTLD-TDP43 | Female | 69 | 13:25 | *C9orf72* |
| 55 | FTLD-TDP43 | Female | 58 | 11:00 | *C9orf72* |
| 56 | FTLD-TDP43 | Female | 57 | 04:15 | *C9orf72* |
| 57 | FTLD-TDP43 | Male | 55 | 12:00 | *C9orf72* |
| 58 | FTLD-TDP43 | Male | 69 | 05:45 | *C9orf72* |
| 59 | FTLD-TDP43 | Female | 66 | 11:30 | *C9orf72* |
| 60 | PiD | Male | 74 | 07:00 |  |
| 61 | PiD | Male | 56 | 09:30 |  |
| 62 | PiD | Male | 74 | 10:15 |  |
| 63 | PiD | Female | 66 | 13:30 |  |
| 64 | CBD | Male | 58 | 13:00 | *MAPT* (p.P301L) |
| 65 | CBD | Male | 58 | 10:00 | *MAPT* (p.P301L) |
| 66 | CBD | Male | 53 | 16:55 | *MAPT* (p.P301L) |
| 67 | CBD | Male | 72 | 05:50 | *MAPT* (p.P301L) |
| 68 | CBD | Male | 75 | 05:55 | *MAPT* (p.P301L) |
| 69 | CBD | Female | 63 | 07:15 | *MAPT* (p.P301L) |

sAD: Sporadic Alzheimer's dementia; fAD: Familial Alzheimer's dementia; PiD: Pick's disease; CBD: Corticobasal degeneration; *C9orf72* indicates patients with an expansion mutation of the GGGCC hexanucleotide repeat (>30 repeats); PMI: Postmortem interval.

Our evaluation through qPCR using divergent oligonucleotides revealed direction effects that were consistent with those reported by Dube et al. in the context of AD, with *circHOMER1*, *circKCNN2*, *circFMN1* and *circMAN2A1* showing a trend towards a decreased expression, and *circDOCK1*, *circMAP7*, *circMAN2A1*, *circRTN4*, and *circPICALM* presenting a tendency towards an enriched expression in AD patients compared to controls (Figure 1). However, only in the case of *circDOCK1* and *circHOMER1* these differences reached statistical significance in the sporadic AD (sAD) group. Notably, in accordance with Dube *et al.* work, the group of fAD, conformed by subjects with Mendelian forms of disease, presented more striking differences in the levels of circRNAs compared to controls than the non-genetic forms of the disease, with 4/8 circRNAs showing significant differences between fAD and controls.

**Figure 1.**
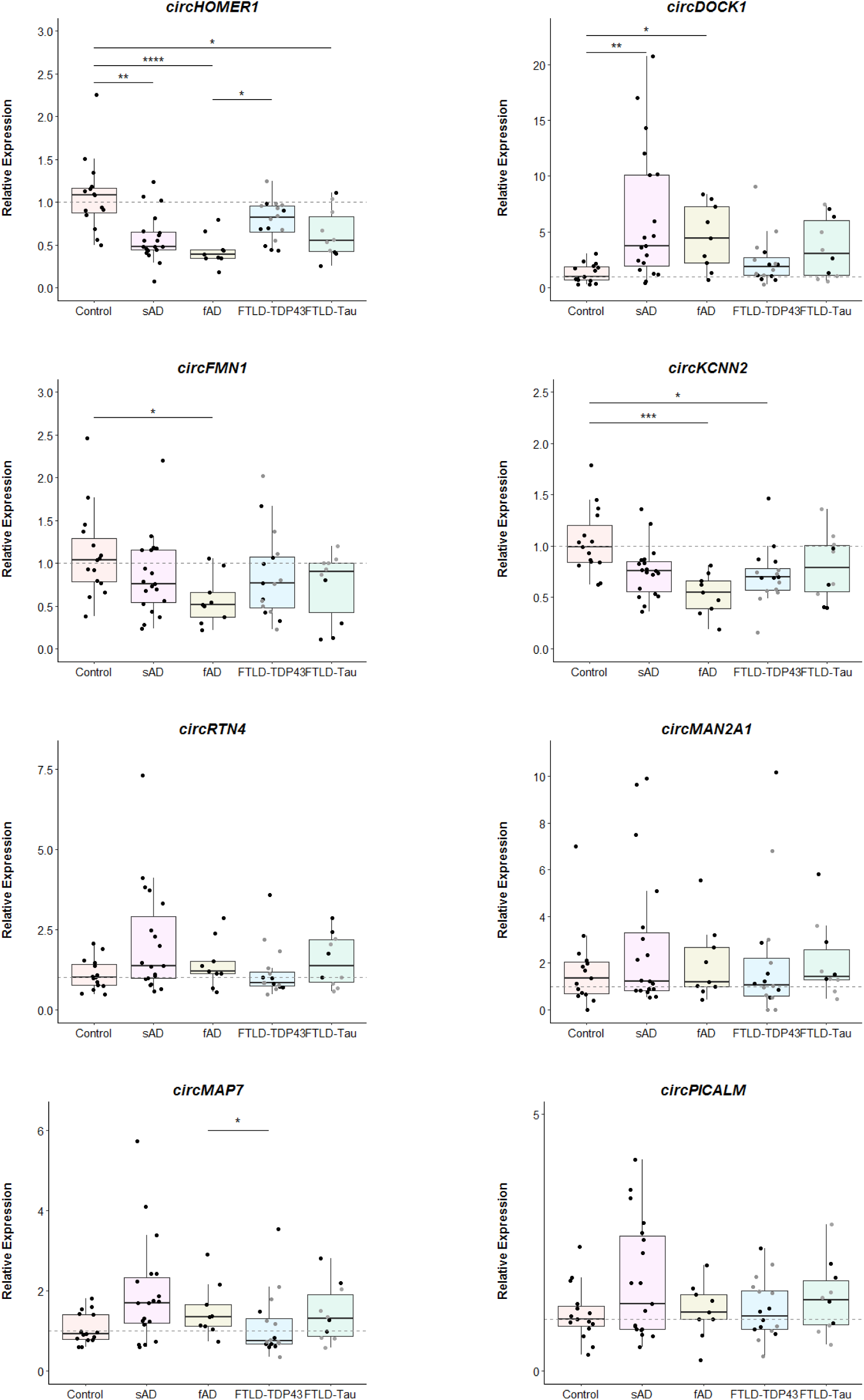
cricRNA expression levels in each individual grouped by the five major phenotypes. Brain samples with pathogenic mutations in the FTLD-Tau and FTLD-TDP43 groups are depicted by a grey circle. * *P* < 0.05; ** *P* < 0.01; \*\*\**P* < 0.001; **** *P* < 0.0001.

In order to assess whether these changes were specific to AD, we evaluated the expression pattern of these circRNAs in FTLD. The levels of *circKCNN2* were significantly reduced in the FTLD-TDP43 group compared to healthy controls. Among FTLD-Tau, levels of *circHOMER1* were also decreased compared to controls. When genetic causes of FTLD due to mutations in *C9ORF72* or *MAPT* were analyzed independently, we did not find any significant change in the expression of any circRNAs compared to controls (data not shown). Also, no correlation of circRNA levels with the age of death was found in the whole sample, or when the analysis was stratified by the 5 phenotype groups (data not shown).

## Discussion

Overall, our results reinforce and complement the findings by Dube and collaborators, showing that the levels of some circRNAs are altered in brain tissue of AD patients. Interestingly, our results in the autosomal-dominant AD group revealed even larger changes in circRNA compared to the sporadic counterpart, confirming previous results. Our correlation analyses partially precluded a possible age-related effect that could influence these outcomes. However, it is important to note that our fAD series was significantly younger than controls. Perfectly age-matched controls with younger age at death to compare circRNA levels with the fAD group would be an invaluable resource to better understand these substantial differences in this particular form of disease. The inclusion of patients with FTLD in our study revealed that some significant changes in circRNA levels can also be observed in other neurodegenerative disorders. A good example is the significant decrease of the expression levels of *circHOMER1* in FTLD-Tau. The presence of *circHOMER1* in neuronal cell bodies and dendrites, its substantial expression in the hippocampus (a region particularly vulnerable in AD and other dementias), and the alterations of its expression levels due to neuronal activity^1^, strengthens its role (and maybe the role of other circRNAs with similar behaviors) across different neurodegenerative disorders.

Although recently discovered, these heterogeneous groups of transcripts are very promising candidates to study in order to better understand the biological bases of neurodegenerative disorders, discover novel peripheral biomarkers and may give rise to novel therapeutic targets.

## Acknowledgements

This work was supported in part by Carlos III Institute of Health, Spain (grant number PI18/00326), Fondo Europeo de Desarrollo Regional (FEDER), Unión Europea, “Una manera de hacer Europa”. We are also indebted to the Neurological Tissue Bank of the Biobanc-Hospital Clínic-IDIBAPS (Barcelona) for providing human brain samples.

